# Evasion of neutrophil-mediated bacterial clearance in *Pseudomonas aeruginosa* isolates from new-onset infections in cystic fibrosis children

**DOI:** 10.1101/2024.09.29.615549

**Authors:** Kelly Kwong, Sophia Goldman, Annie Beauchamp, Karim Malet, Ines Levade, Lucia Grana, David S. Guttman, Valerie Waters, Dao Nguyen

## Abstract

Chronic *Pseudomonas aeruginosa* (PA) infections in cystic fibrosis (CF) patients can persist for decades and are associated with poor clinical outcomes. New-onset PA infections are routinely treated with antibiotics, but unfortunately up to 40% of patients fail eradication therapy due to reasons that are poorly understood. Recently, we found that Persistent PA isolates from CF patients who failed tobramycin eradication therapy were more resistant to *in vitro* neutrophil-mediated opsonophagocytosis and intracellular bacterial killing (OPK) and were significantly associated with a non-twitching phenotype compared to Eradicated isolates. In this study, we sought to investigate how Persistent isolates evade in neutrophil-mediated bacterial clearance *in vitro* and whether these PA isolates also persist *in vivo*. Furthermore, we investigated whether restoring pilus-mediated twitching motility is sufficient to restore susceptibility to *in vitro* OPK and *in vivo* bacterial clearance. Using primary murine serum and bone marrow-derived neutrophils, we demonstrated that Persistent isolates are resistant to several neutrophil antibacterial functions compared to Eradicated isolates. Additionally, mice failed to clear pulmonary infections caused by Persistent isolates but not Eradicated isolates despite comparable responses in leukocyte recruitment and cytokine responses. We demonstrate that loss of Type IV pilus-mediated twitching motility confers a fitness advantage for a Persistent isolate during a murine pulmonary infection, and restoration of pilus-mediated twitching motility improves *in vivo* bacterial clearance. Our findings show that resistance to neutrophil-mediated bacterial clearance in Persistent isolates are partly mediated by loss of Type IV pilus-dependent motility and contributes to the persistence of new onset PA infections.

## Introduction

Cystic fibrosis (CF) is a multi-system genetic disease caused by mutations in the cystic fibrosis transmembrane conductance regulator (CFTR) gene. CF lung disease, the major most cause of morbidity and death. is characterized by a vicious cycle of persistent infection and sustained inflammation leading to chronic respiratory symptoms, acute worsening of symptoms in events termed pulmonary exacerbations, and progressive decline in lung function [1,2]. The Gram-negative opportunistic pathogen *Pseudomonas aeruginosa* (PA), causes chronic airway infections in up to 60% of adults with CF and is associated with worse clinical outcomes [3–6]. Antibiotics, such as inhaled tobramycin, are routinely used to treat new-onset PA infections in an attempt to eradicate PA infections at an early stage and prevent progression to chronic infections [7–10]. Unfortunately, eradication therapies fail in up to 40% of patients even in the absence of antibiotic resistance, resulting in PA infections that persist [11], and the reasons for failed PA eradication remain incompletely understood. Notably, no clinical predictors have been found to be significantly associated with failed eradication [11–13].

Neutrophils are the primary effector immune cells required for clearance of PA *in vivo*, and their function can enhance antibiotic eradication therapy. Phagocytosis and intracellular bacterial killing through ROS-dependent and independent mechanisms are hallmarks of neutrophil antimicrobial functions [14]. However, numerous PA factors can modulate these complex processes and impede host-mediated clearance [15,16]. Mayer Hamblett et al. [17] and Vidya et al. [11] previously characterized the phenotypes of PA isolates infecting CF children from two clinical studies of antibiotic eradication therapy for new-onset PA and showed that strain-specific phenotypes such as mucoidy, lack of twitching motility and wrinkly colony morphology (overproduction of Psl and Pel exopolysaccharides) were associated with persistent infection following antibiotic eradication therapy. These findings led us to characterize the collection of PA isolates collected from CF children from Vidya et al. [11] using *in vitro* neutrophil opsonophagocytic killing (OPK) assays to examine the link between specific PA phenotypes, their resistance to neutrophil antibacterial functions, and outcomes of eradication therapy in CF patients [18]. We showed that infection with PA isolates resistant to *in vitro* neutrophil phagocytosis was a significant independent predictor of failed eradication therapy. More specifically, PA isolates from persistent infections (referred to as Persistent isolates) were more resistant to neutrophil-mediated phagocytosis and intracellular bacterial killing compared to those from eradicated infections (referred to as Eradicated isolates). Deficiency in twitching (i.e., the type IV pili-mediated surface motility), and mucoidy (i.e., alginate overproduction) were the bacterial phenotypes most strongly associated with impaired *in vitro* neutrophil antibacterial functions [18].

In this study, we sought to understand how Persistent isolates evade neutrophil-mediated bacterial clearance. We chose four representative Persistent and Eradicated isolates and investigated the bacterial-neutrophil interactions *in vitro* and their implications for bacterial clearance *in vivo.* Using primary murine neutrophils, we demonstrated that Persistent isolates were resistant to complement binding on the bacterial surface, displayed reduced bacterial adhesion to neutrophils, elicited lower phagocytosis and neutrophil degranulation and were less susceptible to intracellular killing compared to Eradicated isolates. Furthermore, mice failed to clear pulmonary infections caused by Persistent isolates but not Eradicated isolates, despite comparable responses in leukocyte recruitment and cytokine responses. Notably, functional restoration of Type IV pili-mediated motility with PilA overexpression in a non-piliated non-twitching Persistent isolate restored the isolate’s susceptibility to neutrophils and bacterial clearance *in vivo* in a mixed strain infection. Our data suggests that neutrophil interactions with Persistent isolates are impaired compared to those with Eradicated isolates. This is in part driven by loss of Type IV pilus-mediated twitching motility and contributes to failed *in vivo* clearance.

## Materials and methods

### Study design and PA isolates

PA clinical isolates were recovered from patients’ sputum collected immediately before the initiation of inhaled tobramycin as part of the Early PA Eradication Study and CF Sputum Biobank at the Hospital for Sick Children (REB#1000019444) as previously described [11]. New-onset PA infection was defined as a PA-positive sputum culture with at least 3 PA-negative sputum cultures in the previous 12 months, and a sputum culture was collected one-week after tobramycin treatment to determine the outcome of eradication therapy [19,20]. PA infection was considered “persistent” if the post-treatment sputum culture was positive for PA, and “eradicated” if the post-treatment sputum culture was negative. For this study, we selected two pairs of representative clinical PA isolates from persistent and eradicated infections, referred to as Persistent and Eradicated isolates respectively, based on their susceptibility to *in vitro* neutrophil phagocytosis and intracellular bacterial killing as previously determined [18].

### Bacterial culture and phenotypic characterization

To generate bacterial suspensions for all assays described below, PA isolates were grown in 5 mL Luria-Bertain (LB) broth (Thermo Scientific) overnight at 37 °C with shaking at 200 rpm, spun down for 5 min at 12,000 rpm, washed once and resuspended in sterile phosphate buffered saline (PBS) to the indicated optical density at 600nm (OD_600_) as specified in each assay. Assays for twitching and swimming motility, biofilm formation, Congo red binding of bacterial aggregation, and mucoidy status were performed as previously described [18]. Briefly, twitching motility was determined by inoculating a single colony into 1% thin LB agar plate for 2 days at 37 °C and measuring the diameter of the twitching zone after staining with 0.1% crystal violet. Swimming motility was determined by inoculating a single colony into 0.3% LB agar and measuring the diameter of the zone of bacterial growth following overnight incubation at 37 °C. Biofilm formation was measured by inoculating 96-well polystyrene plates with 100 μL of diluted overnight bacterial cultures in LB medium, and incubating overnight under the static condition at 37 °C. The adherent biofilm biomass was stained with 0.1% crystal violet, resolubilized with 95% ethanol, and measured for absorbance at OD_600_. Mucoidy status was determined by colony morphology following growth on yeast extract mannitol (YEM) agar for 24-48 h. Lastly, exopolysaccharide (EPS)-mediated bacterial aggregation was assessed with the colorimetric Congo red binding assay and rugose colony morphology on Congo red agar plate [21].

### Psl surface expression

Psl expression was measured by ELISA as done in [22] with minor modifications. Bacteria grown overnight and resuspended in PBS (OD_600_ 0.2) were used to coat Nunc MaxiSorp^TM^ plates (Thermo Scientific 44-2404-21). Following overnight incubation at 4°C, the plates were centrifuged at 7,000 rpm for 5 min, and the supernatants were discarded. Primary anti-Psl antibody Psl0096 (AstraZeneca) was added (1:12000) to PBS containing 1% BSA and 0.1% Tween-20 for 1.5 h at room temperature (RT), followed by 1:25000 donkey anti-human HRP-conjugated secondary antibody (Jackson Immuno Research Laboratories 709-035-149) for 30 min and detection with TMB SureBlue peroxidase substrate (BD Biosciences 555214) was measured at OD_450nm_.

### Congo red binding of Psl and Pel-mediated aggregation on solid medium

Congo red binding of Psl and Pel on solid medium was performed as previously described [21]. Briefly, 2 µL of overnight bacterial culture was spotted on the surface of Vogel-Bonner minimal medium agar containing Congo red for 18 h at 37 ^ο^C. Wrinkly colony morphology was assessed in a blinded fashion by two readers.

### C3 complement deposition

C3 binding to bacterial surfaces was measured by flow cytometry. Bacterial suspensions (10^9^ cells/mL in sterile PBS) were opsonized with 20% pooled mouse serum (C57BL/6 mice, Charles River Laboratories) for 1 h at 37 °C. Samples were then washed with PBS, stained with 1:500 primary C3 polyclonal antibody (Thermo Scientific PA5-21349) for 1 h on ice, followed by staining with 1:500 goat anti-mouse Alexa Fluor 647 secondary antibody (Thermo Scientific A-21244) for another 1 h. 10,000 events were acquired for each sample using the Fortessa flow cytometer (BD Biosciences) with gating on the AF647+ population and using non-opsonized bacteria as negative controls after gating to exclude debris and doublets.

### Primary murine neutrophil isolation

For all *in vitro* neutrophil assays, bone marrow-derived neutrophils (BMDN) were isolated from C57BL/6 male mice (8-10 weeks, Charles River Laboratories) using STEMCELL EasySep^TM^ mouse neutrophil negative selection kit (STEMCELL 19762) according to the manufacturer’s instructions.

### Neutrophil opsonophagocytosis and intracellular bacteria killing

Assays to measure *in vitro* phagocytosis and intracellular bacterial killing were performed as previously described using BMDN [18]. Bacterial suspensions (10^7^ cells/mL in Hank’s balanced salt solution without calcium, magnesium, and phenol red (HBSS) (Wisent 311-511-CL)) were opsonized with 10% pooled mouse serum, followed by incubation with 2.5 x 10^5^ neutrophils at a multiplicity of infection (MOI) of 10 for 30 min at 37 °C. Gentamicin 100 μg/mL was then added for 30 min (T=60 min for phagocytosis) or 1.5 h (T=120 h for intracellular bacterial killing) to kill extracellular PA. BMDN were then washed with HBSS, lysed with 0.1% Triton X-100 and plated on LB agar to measure viable intracellular bacteria by colony-forming unit (CFU) counts. For our calculations, the input bacterial is the total bacteria at T = 0 min (initial inoculum). Phagocytosis index was calculated as the number of internalized bacteria at T = 60 min per 5×10^5^ input bacteria. The intracellular bacterial killing was calculated as the relative reduction in intracellular bacterial burden, namely [the number of internalized bacteria at 60 min minus the number of internalized bacteria at 120 min] per 5×10^5^ input bacteria.

### Bacterial adhesion to neutrophils

Bacterial suspensions were prepared at 1.2 x 10^7^ cells/mL with HBSS. 2.5 x 10^5^ BMDN per well were seeded on a microscopic multi-chamber slide (Labtek 154534) followed by incubation with cytochalasin B (Sigma C6762) for 30 min at 37 °C to inhibit phagocytosis. BMDN were then incubated with PA at MOI 20 in the presence of 10% pooled mouse serum for 1 h at 37 °C. Following infection, BMDN were washed with PBS and blocked with 10% BSA overnight at 4 °C. Primary anti-PA antibody (1:1000) (Abcam ab68538) was used to stain for extracellular PA at 4 °C overnight followed by staining with goat anti-mouse Alexa Fluor 647 secondary antibody (Thermo Scientific A-21244) (1:500) for 1 h at room temperature. All samples (infected and uninfected BMDNs) were also stained with DAPI (Thermo Scientific D1306) in the presence of 1% Triton-X100 for 1 min at RT and washed with PBS three times prior to confocal microscopy imaging. BMDN were visualized by confocal microscopy (Zeiss LSM 700) using a 63x objective. Approximately 3 to 4 fields of views per experiment from 2 independent experiments were acquired, and 200 cells were analyzed using the Icy automated cell quantification software [23]. The proportion of bacteria associated with neutrophils was calculated as the number of neutrophils with relative fluorescence intensity (RFI) higher than uninfected neutrophils over the total number of neutrophils.

### Neutrophil CD63 surface expression

We measured surface CD63 expression on BMDN as a marker of neutrophil degranulation following infection with PA. Bacterial suspensions (10^7^ CFU/mL in HBSS) were opsonized with 10% pooled mouse serum and incubated with 5 x 10^5^ neutrophils (MOI 10) for 2 h at 37 °C. Neutrophils were then washed, fixed, and stained with viability marker eFluor 780 (1:1000) (Thermo Scientific 65-0865-14), followed by staining with 1:100 of APC-conjugated anti-murine CD11b, BV510-conjugated anti-murine Ly6G and PECy7-conjugated anti-murine CD63 (Biolegend). For each sample, 100,000 events were acquired by Fortessa flow cytometer (BD Biosciences). Gating was performed to exclude debris and doublets, live CD63 positive neutrophils were identified as eFluor 780^-^, CD11b-APC^+^, Ly6G-BV510^+^, and CD63-PECy7^+^.

### Neutrophil intracellular oxidative burst

We used the Luminol assay to measure reactive oxygen species (ROS) production in neutrophils as previously described [24]. Bacterial suspension (5×10^8^ cells/mL in HBSS with calcium and magnesium) were opsonized with 10% pooled mouse serum for 30 min at 37 °C, then washed, resuspended in HBSS (with calcium and magnesium) and incubated with neutrophils (MOI 50) in the presence of 50 μM of Luminol (Sigma 123072), 2000 U of catalase (Sigma C-3155), and 50 U of superoxide dismutase (Sigma S-7571) in white microtiter plates (PerkinElmer) at 37 °C. ROS generated by neutrophils were measured by luminescence every 5 min using a microplate reader (Tecan Infinite 200, Lifesciences). The area under the curve (AUC) was calculated to determine the relative ROS generated by neutrophils over the time course of 2 h. Results were normalized to ROS induced by phorbol myristate acetate (PMA) to account for variation between experiments (Figure S1).

### PilA complementation

Genetic complementation of *pilA* was performed using a Plac-*pilA* construct that overexpresses *pilA* under control of the synthetic *lac* promoter. The Plac-*pilA* construct was generated in a miniTN7-based vector using the Gibson assembly system (NEB), and chromosomally inserted into strain 565P. The engineered 565P Plac-*pilA* strain (noted *pilA+*) encodes the *aacC1* gene gentamicin cassette and was grown with gentamicin selection as indicated. The primers used - *pilA* were: Forward 5’-ATA GAT CTA. AAC TAT GAC AAT-3’ and reverse 5’-AGC CTT TTT GAG CTT TCA TGC TTA ATT TCT CCT CTT TAA TT-3’ (fragment 1); forward 5’-CGA AAG GTT GTG ATA ACT AAG AAT TCG ACG AGC CTG CTT TT-3’and reverse 5’ATG GTA AGC CCT CCC GTA TCG-3’ (fragment 2); forward 5’-ATA CGG GAG GGC TTA CCA TC-3’ and reverse 5’ATT GTC ATA GTT TAG ATC TAT-3’ (fragment 3).

### Murine PA pulmonary infection

A murine PA pulmonary infection model was used to compare the *in vivo* clearance of different PA isolates. Specific pathogen-free C57BL/6 male mice (8-10 weeks, Charles River Laboratories) were anesthetized and infected with 1×10^7^ CFU/mouse using a non-surgical endotracheal method [25]. At designated time points (6, 48, 96 h) post-infection (p.i.), mice were euthanized, blood was collected by cardiac puncture, and the pulmonary vessels were perfused with ice-cold PBS. Serum was collected and stored at −80 °C until analysis.

The bronchoalveolar lavage fluid (BALF) was collected by injecting and aspirating 4 x 500 μL of ice-cold PBS through an endotracheal catheter, and cells were pelleted down (1,500 rpm, 10 min). Aliquots of the BALF cell-free supernatant were stored at −80 °C with protease inhibitor cocktails (Thermo Scientific 78441) until analysis. Following the lavage, lungs were removed, placed in 10 mL ice-cold PBS, minced with razor blades and aliquoted for the following processing steps. To quantify the viable PA bacterial burden, an aliquot of minced lung was homogenized (0.2 mm stainless steel beads, Bullet Blender, Next Advance) serially diluted and plated on Pseudomonas isolation agar (PIA) for colony forming units (CFU) counts. The remaining minced lung sample was split into two parts. For cytokine measurements, an aliquot of minced lungs was homogenized in the bullet blender, pelleted down (12,000 rpm, 5 min), and the cell-free supernatants were stored at −80 °C with protease inhibitor until analysis. For flow cytometry analyses, an aliquot of minced lungs was digested with collagenase (Sigma C5138) and filtered through a 100 μm cell strainer. Following red blood cell lysis, cells were spun down, washed, resuspended in PBS, and immediately processed for flow cytometry.

### Mixed strain infection murine pulmonary infection

For competition experiments with mixed strain infection, bacterial isolates 565P and *pilA+* were streaked onto Luria-Bertain (LB) agar plates with and without gentamicin (50 μg/mL), respectively, and grown overnight at 37°C. Overnight cultures of 565P and *pilA+* containing gentamicin (25 μg/mL) were spun down for 5 mins at 12,000 rpm, washed, resuspended in PBS and diluted to an estimated concentration of 2 x 10^8^ CFU (OD_600_ =0.20), and mixed at a 1:1 ratio. Mice were anesthetized and infected with 1 × 10^7^ total bacteria in 50 μL of the mixed inoculum as done with the single strain infection. At 6 h, 24 h and 96 h p.i, the lungs were collected and processed for viable bacterial counts as follows. Serial dilutions of lung homogenates were plated onto PIA plates with and without gentamicin (Gm 25 μg/mL). Plates were incubated overnight at 37°C, and CFU counts were enumerated for 565P and *pilA+* respectively. The *pilA+* were counted as viable CFU on PIA Gm plates, while the 565P counts were calculated as the total viable CFU on PIA minus the CFU on PIA Gm plates. The competitive index (CI) was calculated as output [*pilA+* / 565P] CFU ratio divided by the input [*pilA+* / 565P] CFU ratio [26–28]. A CI value equal to 1 indicates that both strains persisted similarly and neither had a competitive advantage, while a CI value < 1 indicates that 565P outcompeted *pilA+*.

All animal experiments were carried out in accordance with the Canadian Council on Animals Care and with approval from the Animal Care Committee of the Research Institute of the McGill University Health Centre (AUP #7586).

### Cytokine and chemokine measurements

Serum, protease-inhibitor treated BALF and lung homogenate supernatant samples were analyzed for a panel of mouse cytokine/ chemokine using Luminex 31-Plex Discovery Assay Array (MD31) with the following biomarkers: Eotaxin, G-CSF, GM-CSF, IFNγ, IL-1α, IL-1β, IL-2, IL-3, IL-4, IL-5, IL-6, IL-7, IL-9, IL-10, IL-12p40, IL-12p70, IL-13, IL-15, IL-17A, IP-10, KC, LIF, LIX, MCP-1, M-CSF, MIG, MIP-1α, MIP-1β, MIP-2, RANTES, TNFα, VEGF-A (Eve Technologies).

### Flow cytometry analysis of BALF and whole lung homogenates

Two million cells from whole lung single-cell suspensions or BALF were fixed and stained with eFluor780 fixable viability dye (1:1000) (Thermo Scientific 65-0865-14), then surface stained with fluorescently-conjugated murine innate immune cell markers that target neutrophils (CD45-PE, CD11b-APC, Ly6G-BV510), monocytes (CD45-PE, CD11b-APC, Ly6C-BV785), macrophages (CD45-PE, CD11b-APC, Ly6G^lo^-BV510, F4/80^hi^-Pacific blue) and dendritic cells (CD45-PE, CD11b-APC, Ly6G^lo^-BV510, CD11C-BV711). 100,000 events were analyzed by flow cytometry (Fortessa, BD Biosciences) with gating on live CD45+ cells after excluding debris and doublets. Unless specified, all antibodies were from Biolegend and used at a final concentration of 1:100. Flow cytometry data were analyzed using FlowJo Software, version 10.8.1 (Ashland, OR, USA).

### Whole genome sequencing and analysis

All PA isolates were sequenced using Illumina NextSeq and analyzed as described in [29]. Sequencing reads were previously deposited in NCBI (PRJNA556419). As previously described in [30], following de-novo assembly, sequencing read quality assessment and adaptor trimming, targeted comparative genome analysis was then performed on the two pairs of representative Persistent (565P and 505P) and Eradicated (513E, 558E) isolates by using Trimmomatic [31] and Spades [32] v3.14.1, respectively. Annotation was performed with Prokka [33].

### Statistical analyses

All results are shown as mean ± SEM or median ± IQR as indicated. Statistical analyses were done using Prism 9 software (Graphpad). Comparisons were performed using ANOVA Dunn’s test or unpaired two-tailed student’s t-test as indicated. A P value of < 0.05 was considered statistically significant. \**P* < 0.05, \*\**P* < 0.01, and \*\*\**P* < 0.001.

## Results

### Representative Persistent and Eradicated PA isolates

Our previous studies indicated that Persistent isolates were resistant to *in vitro* OPK by dHL-60 cells, had reduced pilus-mediated twitching motility, were more commonly mucoid, and had increased Psl production compared to Eradicated isolates [11,18,57]. We thus chose two pairs of representative Persistent and Eradicated PA isolates from our collection to further investigate bacterial-neutrophil interactions and their implications to bacterial clearance *in vivo*. The isolates were selected based on their susceptibility to neutrophil OPK in our *in vitro* assays, as shown in Figure 1, to represent the median and high/low range of phagocytosis and intracellular bacterial killing. Persistent isolates were also selected to overproduce Psl and Pel (wrinkly colony morphology and high Congo red binding) and be deficient in pilus-mediated twitching motility, while Eradicated isolates did not overproduce Psl and Pel and had functional twitching motility. All the isolates were chosen to be non-mucoid to avoid the confounding effect of mucoidy. The phenotypic characteristics of these isolates are shown in Table 1.

**Figure 1.**
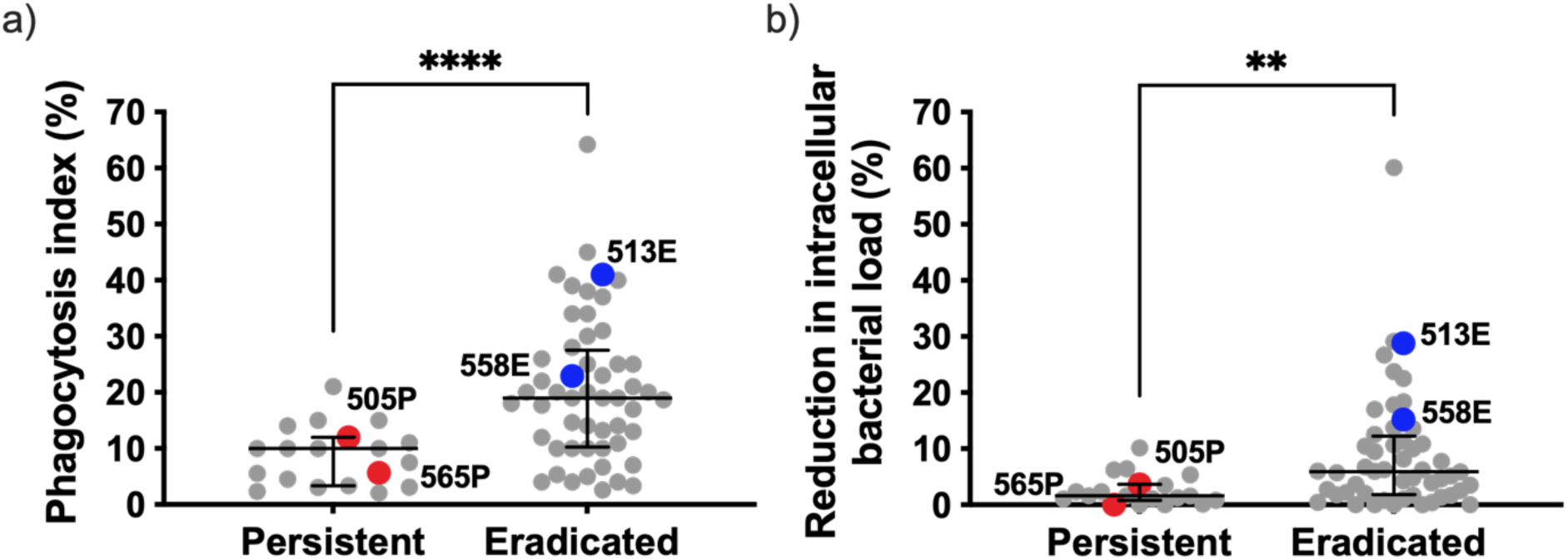
PA clinical isolates from patients with persistent infections are resistant to *in vitro* a) phagocytosis and b) intracellular bacterial killing compared to those from patients with eradicated infections. This figure includes data adapted from Kwong et al. 2022 [18]. Assays were performed with immortalized human neutrophil-like dHL-60 cells. Representative Persistent and Eradicated isolates used in this study are highlighted. Results are shown as median ± IQR (n ≥ 4 biological replicates from n ≥ 2 independent experiments). \*\**P* < 0.01, \*\*\*\**P* < 0.0001 using the Mann-Whitney test.

**Table 1.** Phenotypic characteristics of PA isolates.

| Isolates | Eradication status | Twitching motility (mm) | Swimming motility (mm) | Biofilm formation (OD<sub>600nm</sub>) | Mucoidy | Congo Red binding | Wrinkly colony morphology | Psl production (OD<sub>450nm</sub>) |
| --- | --- | --- | --- | --- | --- | --- | --- | --- |
| **565P** | Persistent | 0 | 0 | 0.12 | - | -0.38 | +++ | 1.8 |
| **505P** | Persistent | 1 | 8 | 0.24 | - | -0.32 | + | 1.9 |
| **513E** | Eradicated | 23 | 13 | 0.18 | - | -0.16 | - | 0.9 |
| **558E** | Eradicated | 26 | 12 | 0.21 | - | -0.07 | - | 1.7 |

Representative Persistent and Eradicated PA isolates were selected according to their susceptibility to *in vitro* antibacterial functions by dHL-60 cells, twitching motility, and EPS production. Among these measurements, pilus-mediated twitching motility ( $P = **$ ), wrinkly colony morphology ( $P = *$ ), and Psl production ( $P = **$ ) were significantly different in a subset of well-characterized Persistent isolates ( $n = 7$ ) compared to Eradicated isolates ( $n = 8$ ). $**P < 0.01$ and $*P < 0.05$ [11,18,57].

### Opsophagocytosis and intracellular bacterial killing

Since our previous study used immortalized human neutrophil-like dHL-60 cells, we first validated the susceptibility of the four representative PA isolates to phagocytosis and intracellular bacterial killing in primary murine BMDN. As expected, Persistent isolates 565P and 505P were more resistant to phagocytic uptake and intracellular bacterial killing compared to Eradicated isolates 513E and 558E (Figures 2a, b). To further characterize the PA strain-specific interactions with neutrophils, we then compared the Persistent and Eradicated isolates in a series of *in vitro* assays.

**Figure 2.**
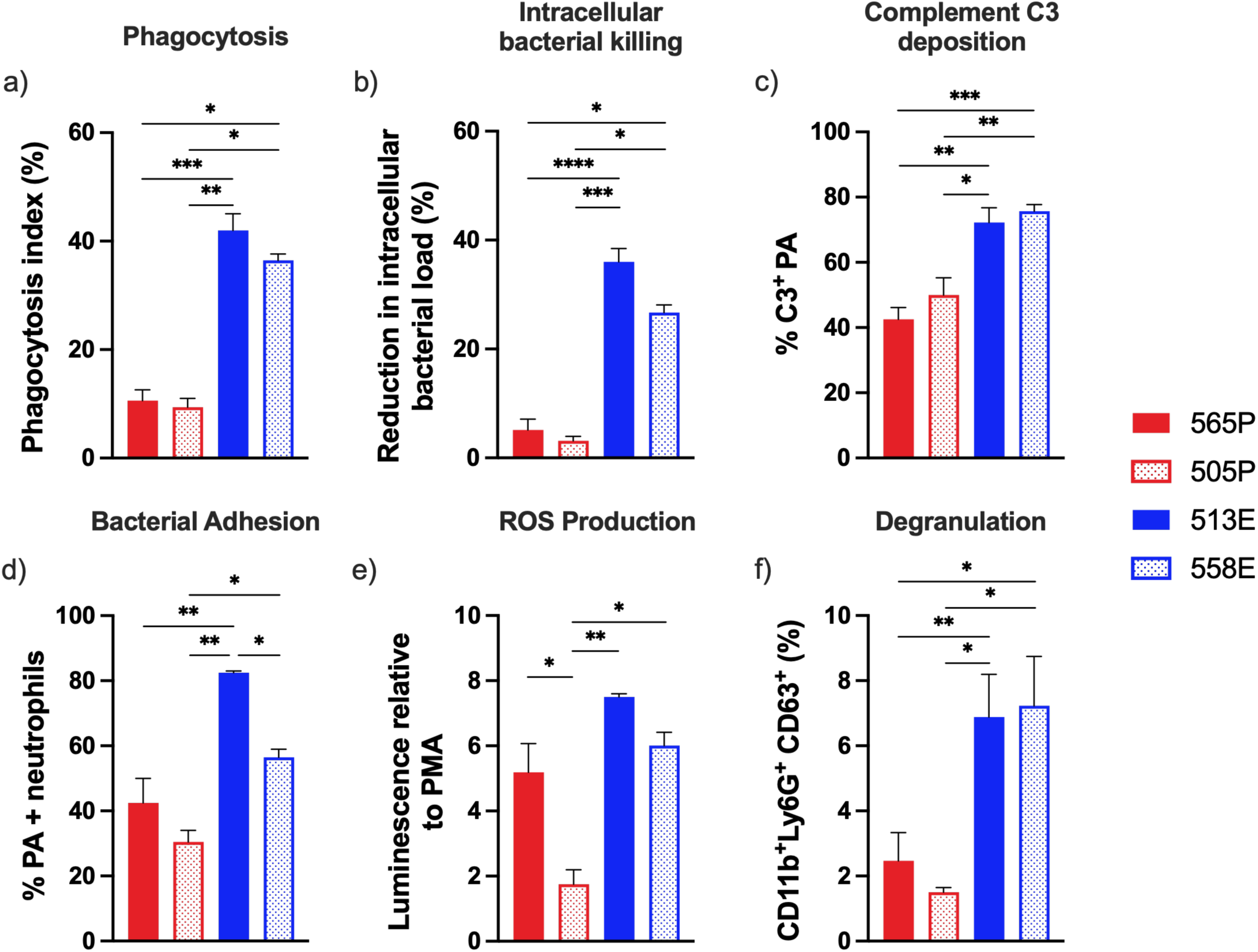
*In vitro* PA-neutrophil interactions with primary murine neutrophils. The Persistent isolates 565P and 505P, and Eradicated isolates 513E and 558E were tested *in vitro* using primary bone derived murine neutrophils for a) Phagocytosis; b) Intracellular bacterial killing; c) C3 complement deposition on PA measured as % C3+ PA cells / total PA; d) Bacterial adhesion to neutrophils measured as % neutrophils associated with PA / total neutrophils in adhesion assay; e) ROS production measured using the luminol assay and reported as luminescence AUC relative to a phorbol myristate acetate (PMA) control; f) Neutrophils degranulation in response to PA isolates measured as % CD11b^+^Ly6G^+^CD63^+^/ total CD11b^+^Ly6G^+^. Results are shown as mean ± SEM (n ≥ 6 biological replicates from n ≥ 2 independent experiments). \**P* < 0.05, \*\**P* < 0.01, \*\*\**P* < 0.001, \*\*\*\**P* < 0.0001 using the ANOVA Dunn’s test for a), b), f), and Tukey’s test for c), d), e).

### Surface binding of complement

Complement-dependent opsonization is important for bacterial clearance, particularly for neutrophil-mediated antibacterial functions [34]. Surface molecules such as the EPS Psl can interfere with complement deposition on the PA bacterial surface [35].

We thus measured the surface binding of C3, the most abundant complement component that binds to the PA surface [35] and observed a significantly lower C3 surface binding on Persistent isolates 565P and 505P (42.6% and 50% respectively) compared to the Eradicated isolates 513E and 558E (72.2% and 75.7% respectively) (Figure 2c).

### Bacterial surface adhesion to neutrophils

Bacterial adhesion to neutrophils is a critical initial step to initiate phagocytic bacterial uptake and be modulated by strain-specific variation in surface deposition of complements, bacterial motility and surface appendages [35,36]. We thus compared the ability of Persistent and Eradicated isolates to bind neutrophils in a bacterial adhesion assay using BMDN. As shown in Figure 2d, a lower proportion of PA-adherent BMDN were observed with 565P or 505P infections compared to 513E or 558E infections (42.5% and 30.5% vs. 82.5% and 56.5%, respectively) (Figure 2d), suggesting that the Persistent isolates had reduced adhesion to neutrophils.

### Neutrophil ROS production and degranulation

To further examine whether the resistance of Persistent isolates to neutrophil intracellular bacterial killing is attributable to impaired oxidative or non-oxidative killing mechanisms, we assessed the intracellular ROS production and degranulation of BMDN infected with different PA isolates. Isolate 505P induced a significantly lower ROS production than 513E and 558E, while 565P did not (Figure 2e), suggesting that impaired ROS production was not sufficient to explain impaired intracellular killing in both Persistent isolates. Next, we assessed neutrophil degranulation as a proxy for non-oxidative antimicrobial killing activity by measuring neutrophil surface CD63 expression in BMDN infected with different PA isolates. As shown in Figure 2f, the proportion of CD63+ neutrophils was significantly lower among 565P and 505P-infected neutrophils compared to those infected with 513E and 558E (Figure 2f). These results suggest that impaired neutrophil degranulation in response to 565P and 505P infection may have contributed to the reduced killing of these Persistent isolates compared to Eradicated isolates.

### *In vivo* bacterial clearance Persistent and Eradicated PA

In order to determine whether *in vivo* clearance of 565P and 505P was also impaired compared to 513E and 558E, we used a murine pulmonary infection model (Figure 3). We quantified the lung bacterial burden at 6, 48 and 96 h p.i and observed no differences in bacterial burden between the different isolates at 6 h p.i. While we observed a decrease in bacterial burden for all isolates over time, bacterial clearance was of 513E was significantly greater compared to 565P (Figure 3a) at 48 h p.i. (median CFU counts 5.5×10^3^ (513E) compared to 5.8×10^4^ (565P)). By 96 h p.i., 513E was completely cleared in all mice and 558E infected mice had a median CFU below the limit of detection, while 565P and 505P infected mice still had a median of 1.1×10^4^ and 3.7×10^3^ CFU/lung respectively and none had completely cleared their infection (Figures 3a, b).

**Figure 3.**
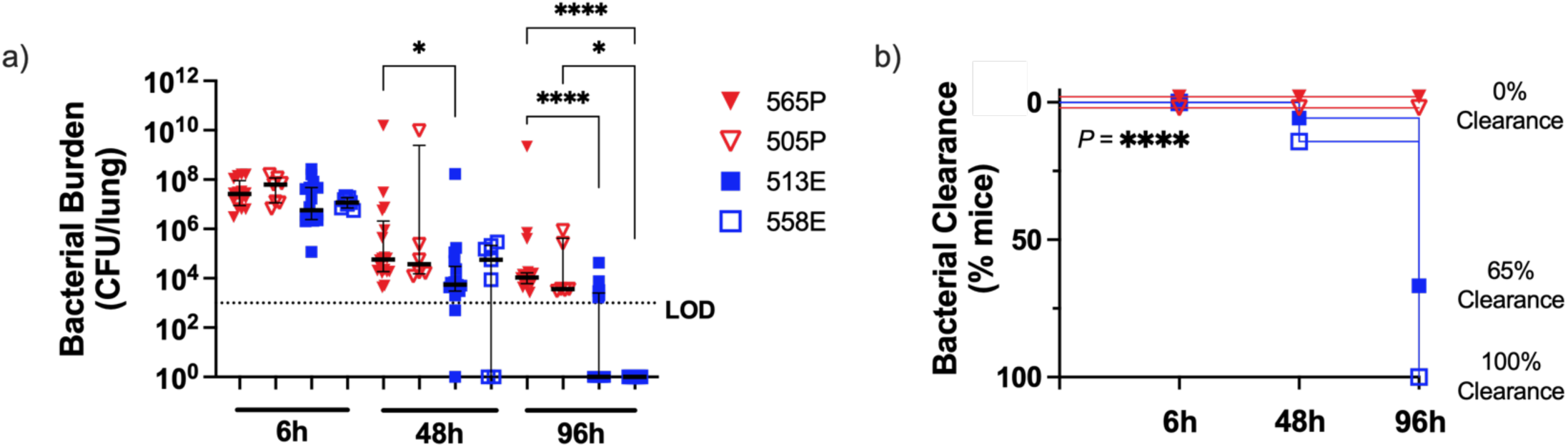
Bacterial clearance of Persistent isolates 565P and 505P is impaired in a murine lung infection model. a) Bacterial burden recovered from whole lung homogenates at 6, 48, and 96 h p.i. Results are shown as median ± IQR. N ≥ 6 mice per strain pooled from ≥ 2 independent experiments. LOD: limit of detection. b) The percentage of bacterial clearance per strain in at 6, 48 and 96 h p.i. \**P* < 0.05, \*\*\*\**P* < 0.0001 using the Kruskal-Wallis test for a) and the log rank test comparing 565P and 505P vs. 558 and 513 infected mice for b).

### *In vivo* immune cell recruitment and cytokine response to Persistent and Eradicated PA infections

To assess whether impaired bacterial clearance of Persistent isolates was due to differential leukocyte recruitment, we measured the immune cell counts in the lung homogenates of mice infected with the different PA isolates at 6, 48, and 96 h p.i. and found no significant differences in the total leukocyte counts (Figure 4a), neutrophil counts (Figure 4b) nor the percentage of neutrophils (Figure 4c) across the different infection groups. Additionally, no significant differences in monocyte, macrophage or dendritic cell counts nor the percentage of these cells was found across the groups (Figure S2). Our data thus far indicates that bacterial persistence in mice infected with 565P or 505P could not be explained by a significant defect in immune cell recruitment.

**Figure 4.**
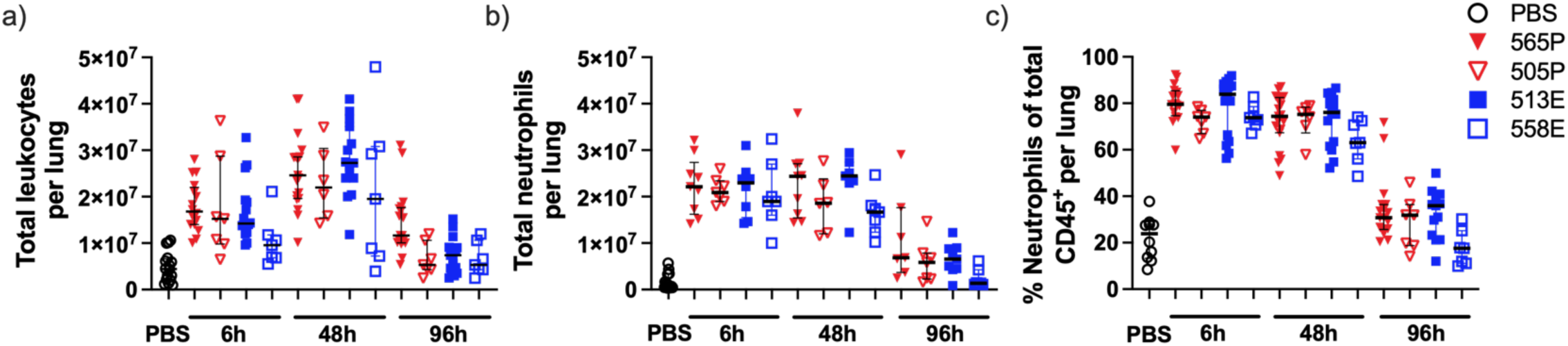
Leukocyte recruitment to the lung in Persistent vs. Eradicated isolate infections. a) Total leukocytes (CD45^+^) in whole lung homogenates, b) Number of neutrophils (CD11b^+^Ly6G^+^CD63^+^), and c) Proportion of neutrophils at 6, 48, and 96 h p.i. Results are shown as median ± IQR. N ≥ 7 mice per strain from ≥ 2 independent experiments. No statistical differences were detected between the groups using ANOVA Dunn’s test.

Next, we measured the host cytokine responses using a 31-plex panel of murine cytokines and chemokines involved in inflammatory and immune responses in serum, lungs, and BALF. Eight key analytes were induced upon PA infection at 6 h p.i., namely the mediators of innate immune cell activation, proliferation, and migration important to bacterial clearance (GM-CSF, G-CSF, MCP-1, IL-6, KC, MIP-1α, IL-1β and TNF-α) [37–41] (Figures 5a-c), consistent with the existing literature [42]. Among these analytes, we noted a significantly lower IL-1β and IL-6 in BALF samples of 505P-infected mice compared to all other ones at 6 h p.i. However, these differences were only observed in 505P-infected mice and nor were they accompanied by reduced immune cell counts, suggesting that they were unlikely to drive the *in vivo* clearance defect observed with both 565P and 505P infections. The other 23 analytes were also analyzed, but their concentrations were below their respective clinical cutoff levels [43] (data not shown). Together, these cytokine profiles do not explain the impaired *in vivo* clearance of both Persistent isolates 565P and 505P.

**Figure 5.**
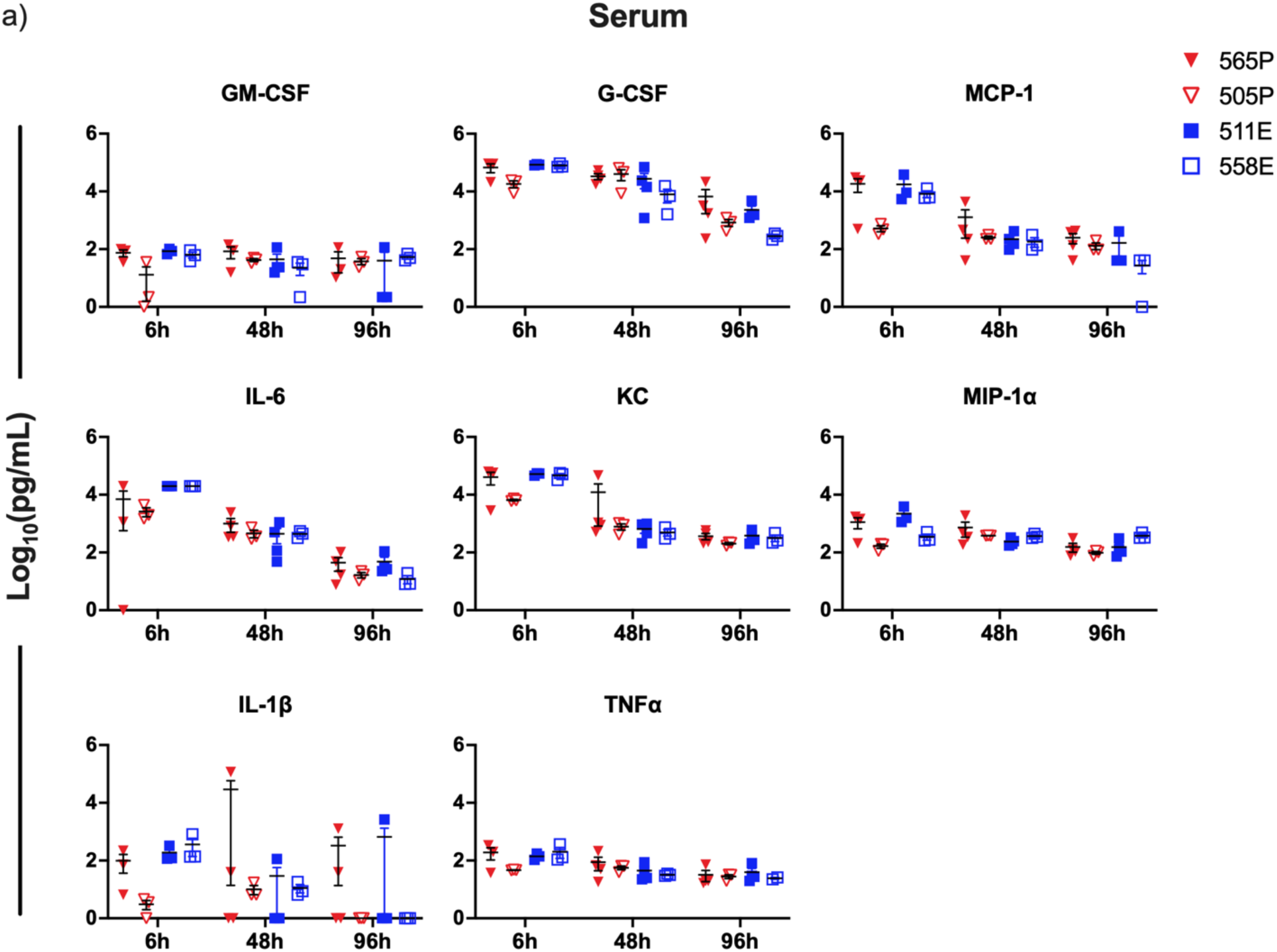
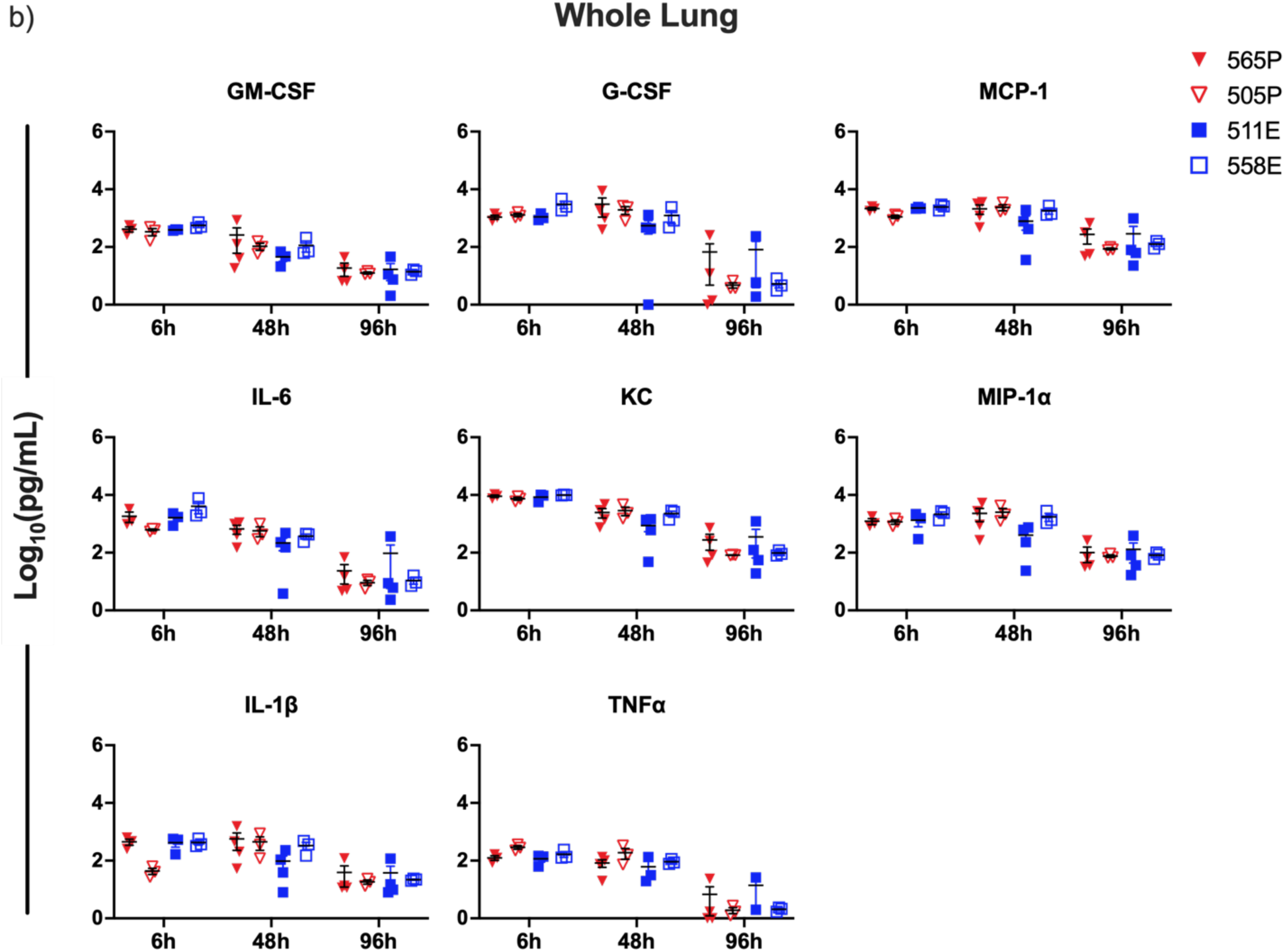
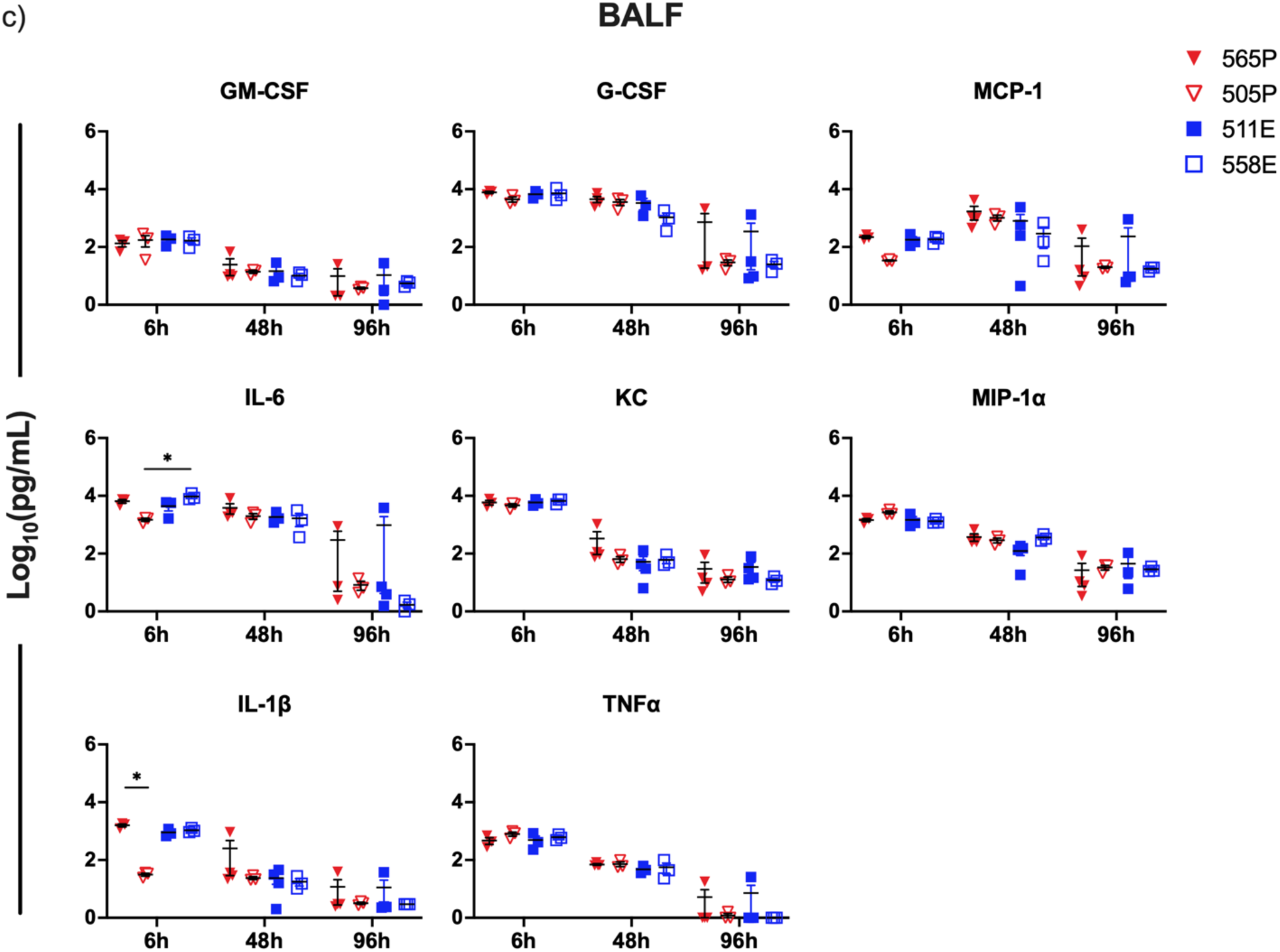
Systemic and lung cytokine responses in mice infected with 565P, 505P, 513E, and 558E isolates. a) Cytokines and chemokines detected in serum, b) Whole lung homogenates and c) the BALF at 6, 48, and 96 h p.i. Results are shown as mean ± SEM (N ≥ 3 mice per time point and isolates) with \**P* < 0.05 using ANOVA Dunn’s test. Out of range (OOR<) data points are represented as 0.

### Targeted genome sequence analysis of Persistent isolates

We hypothesized that loss of type IV pilus-mediated twitching motility is an important contributor to OPK resistance and PA clearance. This phenotype was significantly associated with resistance to neutrophil OPK *in vitro* in our previous study of 71 CF clinical isolates, and 26% (5/19) of Persistent isolates were non-twitching [18]. This led us to examine the contribution of twitching deficiency in the OPK resistance of 565P and 505P. We first performed whole genome sequencing the 4 PA isolates to identify the genetic variation(s) potentially responsible for the twitching defect. We analyzed the sequence of the following genes known to be involved directly or indirectly in type 4 pilus (T4P) biosynthesis and twitching motility: *pilA, pilB, fimV, pilT, pilU, pilS, pilR, RpoN, RetS, fliC, algT/U, tadD, tadB, tadA, rcpC, rsm, flp* [44]. We also examined genes indirectly involved in surface attachment and motility, namely EPS production and global regulators: *mucA, mucD, algK, algE, algG, pslA, pslB, pslH, pslI, pslL, pelA, pelB, pelD, pelF, pelG, cdrA, wspA, wspC, wspE* [45]. We found that *pilA* was absent in 565P, while 505P encoded a functional *pilA* variant allele previously found in pediatric CF isolates (Figure S3) [46]. No other non-synonymous genetic variants were identified in any of the other genes.

### Genetic and functional complementation of *pilA* in the non-twitching 565P isolate

Since the 565P isolate lacked *pilA*, the gene encoding the major pilin subunit of the Type IV pilus, we asked whether overexpressing *pilA* in 565P would functionally complement twitching motility and improve neutrophil antibacterial functions. The twitching motility in the *pilA* complemented strain *pilA+* was restored. Although functional twitching motility remained modest (Figure 6a), this was sufficient to restore significant neutrophil phagocytosis, intracellular bacterial killing, and degranulation (Figures 6b-d). Together, these results suggested that twitching deficiency was an important evasion mechanism of neutrophil antibacterial functions in 565P. Attempts to genetically manipulate and transform the 505P with the Plac-*pilA* construct were unsuccessful.

**Figure 6.**
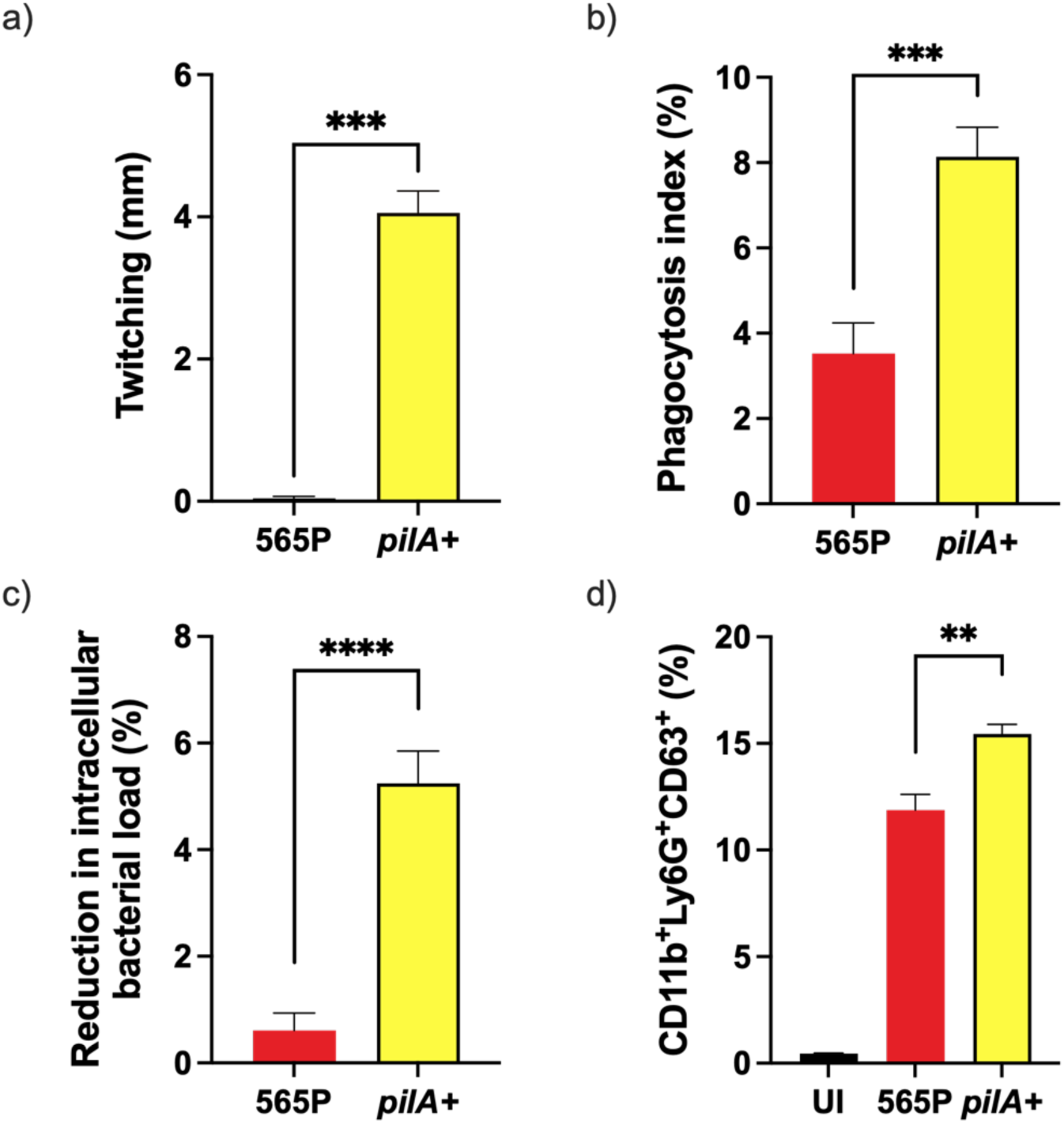
Overexpression of *pilA* restored twitching motility in 565P and its susceptibility to neutrophil antibacterial functions. The 565P strain was complemented by overexpressing Plac-*pilA* in the *pilA+* strain. a) Twitching motility of 565P and *pilA+* as measured by crystal violet staining of the twitching zone; Primary bone marrow derived neutrophils b) Phagocytosis, c) Intracellular bacterial killing, and d) Degranulation in response to *pilA+* compared to its parental 565P strain. Results are shown as mean ± SEM (n ≥ 6 replicates from ≥ 2 independent experiments). \*\**P* < 0.01, \*\*\**P* < 0.001, \*\*\*\**P* < 0.0001 using an unpaired t-test.

### Mixed strain competition *in vivo* and *in vitro*

To further demonstrate whether the loss of pilus-mediated twitching motility in 565P provides a fitness advantage for persistence in the lung by evading neutrophil bacterial clearance, we performed an *in vivo* competition experiment with mixed strain infection. Mice were infected with a 1:1 mixture of 565P and *pilA+*, and each strain can be counted by differential plating in the presence and absence of gentamicin. As shown in Figure 7a, we observed a higher bacterial load of 565P than *pilA+* starting at 6 h (2.6 x 10^6^ CFU/lung vs. 1.2 x 10^6^ CFU/lung) and 24 h p.i. (8.3 x 10^4^ CFU/lung vs. 9.0 x 10^3^ CFU/lung), with a mean competitive index of 0.5 at 6 h and 0.1 at 24 h p.i (Figure 7c). By 96 h p.i, the 565P persisted (3.7 x 10^3^ CFU/lung) while the *pilA+* was eradicated in all but one mouse (median below the CFUlimit of detection) and a mean competitive index of 0.001. Notably, 75% of mice completely cleared *pilA+* from their mixed strain infection by 96 h p.i, while 565P persisted (Figure 7b). These results thus indicate that loss of pilus-mediated twitching motility confers a survival advantage to PA *in vivo*. As controls, we showed that the 565P and *pilA+* strains had comparable growth rates in planktonic cultures. There was also no evidence of inter-strain competition or fitness advantage during mixed strain planktonic laboratory co-cultures (Figure S4).

**Figure 7.**
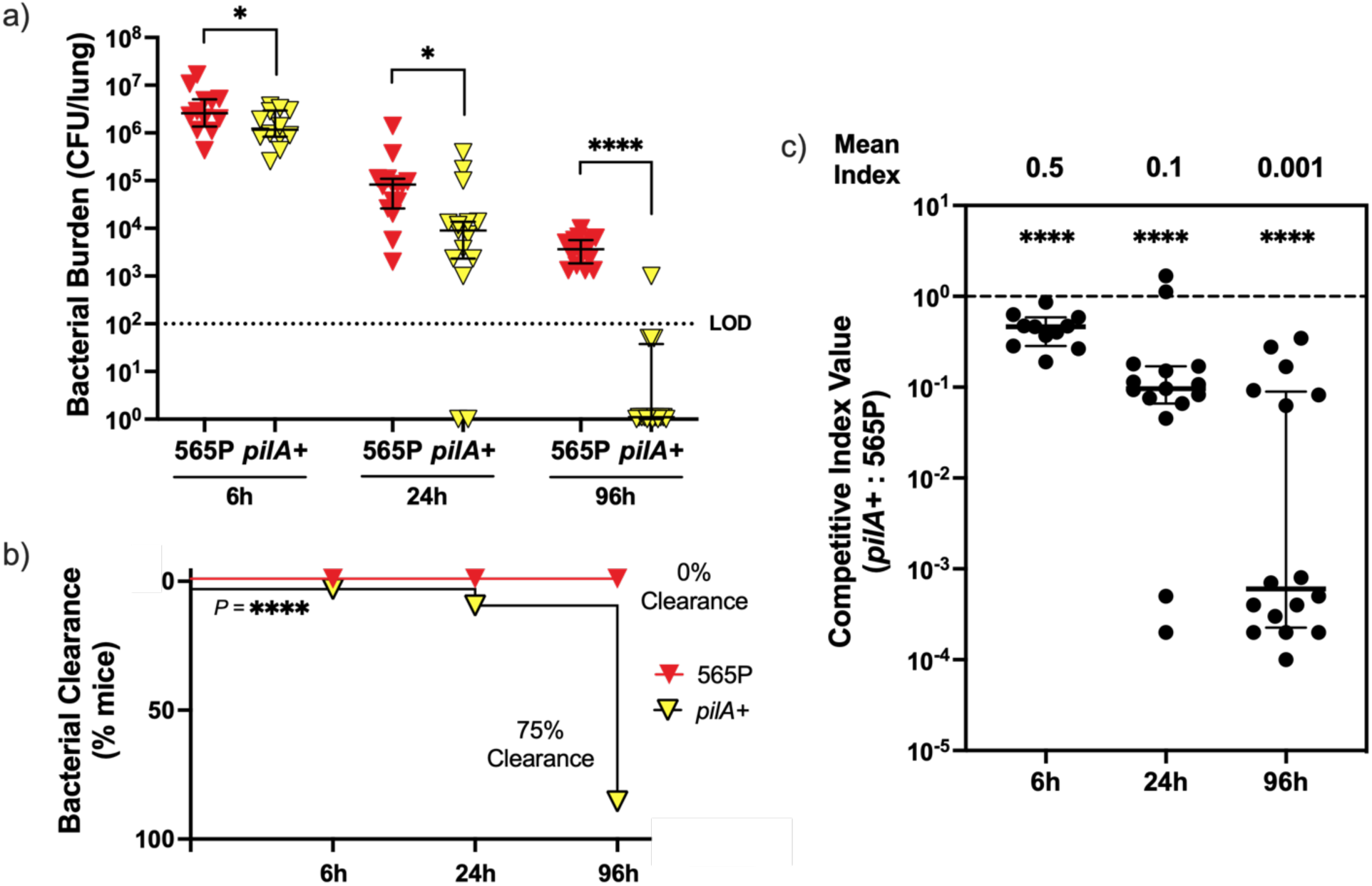
Loss of pilus-mediated twitching motility confers a fitness advantage for Persistent isolate 565P during murine pulmonary infection. a) Lung bacterial burden recovered from 565P and *pilA+* co-infected mice at 6 h, 24 h, and 96 h p.i. Results are shown as median ± IQR. Statistical comparisons were performed using the Mann-Whitney test (\**P* < 0.05, \*\*\*\**P* < 0.0001); LOD = 100 CFU/ mouse; b) Enhanced bacterial clearance was observed in *pilA+* compared to 565P. Statistical comparison was done by the log rank test. N ≥ 11 mice pooled from 3 independent experiments. c) Competitive index analysis of 565P and *pilA+* at 6 h, 24 h, and 96 h p.i. Results are shown as median ± IQR. Asterisks represent a CI statistically different from CI value of 1.0. (\*\*\*\**P* < 0.0001) using the Mann-Whitney test.

## Discussion

Our study sought to understand whether PA strain-specific interactions with neutrophils contribute to the establishment of persistent infection in CF children by testing PA clinical isolates from new onset PA infections in *in vitro* neutrophil assays and a murine infection model. We found that several processes important to neutrophil antibacterial functions (bacterial adhesion to neutrophils, OPK, and CD63 degranulation) were impaired with Persistent isolates 565P and 505P compared to Eradicated isolates 513E and 558E. In contrast to Eradicated isolates, the Persistent isolates also failed to be cleared *in vivo,* and this persistence could not be attributed to impaired cytokine responses or immune cell recruitment. In the non-twitching 565P isolate, restoring its type IV pilus-mediated twitching motility significantly increased its susceptibility to *in vitro* neutrophil OPK and CD63 degranulation activity, and was sufficient to enhance *in vivo* bacterial clearance. Altogether, our findings suggest that strain-specific bacterial phenotypes observed in PA CF clinical isolates, particularly loss of pilus-mediated motility, are important determinants of bacterial-neutrophil interactions that drive bacterial clearance.

Others have previously examined strain-specific PA-neutrophil interactions implicated in bacterial clearance using clinical CF isolates. For example, two studies assessed *in vitro* neutrophil extracellular trap (NET) formation in response to paired clonally related PA isolates from early-stage and chronic CF infections and reported relatively lower NETs elicited by chronic PA isolates compared to their clonally related early isolates [47,48]. Mahenthiralingam et al. originally reported that ∼40% of PA isolates from CF infection lacked swimming (flagellar-mediated) motility and were more resistant to OPK by mouse macrophages compared to motile isolates, but neutrophil phagocytosis was not tested. While they attempted to detect pilus expression, they did not assess twitching (pilus-mediated) motility in these isolates [49].

PA produces three exopolysaccharides (EPS), namely alginate, Psl and Pel, and EPS overproduction contributes to resistance to antibiotic and neutrophil-mediated antimicrobials through several different mechanisms [50]. Overproduction of alginate results in a phenotype called mucoidy, which is common among chronic CF infection isolates [51,52], but relatively rare in environmental and new-onset CF infection isolates [53,54]. Although mucoidy was among the bacterial phenotypes associated with failed eradication therapy [18], and with impaired *in vitro* neutrophil antibacterial functions in our previous study [18], we chose to study representative Persistent and Eradicated isolates that were non-mucoid. Since alginate overproduction is known to inhibit neutrophil elastase activity, NETs, and ROS-mediated antimicrobials [55,56], the study of non-mucoid isolates allowed us to examine the contribution of other bacterial factors without the confounding effect of alginate overproduction.

The EPS Psl and Pel are both cell surface-associated and secreted, and their overproduction leads to a wrinkly colony morphology [45]. Both of our Persistent isolates displayed a wrinkly colony morphology (albeit to a greater degree in 565P), as well as higher Congo red binding, (a surrogate marker for Psl and Pel production [21]), than Eradicated isolates. While our Psl measurement by ELISA did not suggest differences between Persistent and Eradicated isolates, Morris et al. demonstrated that Persistent isolates 565P and 505P expressed high Psl when grown as mature biofilms and measured with immunofluorescence by confocal microscopy using the same antibody [57]. In addition to their role as a constituent of the biofilm matrix, Pel and Psl are important surface molecules that modulate innate immune responses. Psl is involved in cell-substrate bacterial attachment to biotic and abiotic surfaces [58–60], and may thus contribute to the reduced neutrophil adhesion observed in Persistent isolates. Consistent with Mishra et al. who reported that Psl interferes with complement-mediated neutrophil OPK [35], we observed lower deposition of complement C3 on the bacterial surfaces of Persistent compared to Eradicated isolates.

Bacterial evasion of host clearance in murine infection models can be driven by different mechanisms modulating neutrophil-PA interactions and responses. Using a genetically engineered PA mutant *wspF* which overproduces Psl and displays the wrinkly phenotype, Pestrek et al. showed that the *wspF* mutant failed to be eradicated in a mouse acute pulmonary infection model compared to its wild-type parental strain [40]. Notably, the Psl-overproducing *wspF* mutant induced a vigorous neutrophilic inflammatory response *in vitro* and *in vivo*, consistent with observations by Rybtke et al. [61] and ours that neutrophil recruitment is not deficient during infections with Persistent isolates. Interestingly, we did observe a trend towards higher concentrations of IL-1β, KC, IL-6, MCP-1 in serum and BALF from 565P-infected mice compared to 505P-infected mice at 6 h p.i. It is plausible that the increased EPS production seen in 565P increases attachment to host cells and TLR-dependent activation [40]. However, these cytokine differences did not result in detectable differences in immune cell recruitment nor bacterial clearance between those two isolates. Moreover, Baker et al. reported that degradation of Pel increased neutrophil killing of PA biofilms [62], suggesting that Pel may also hinder neutrophil-mediated antibacterial functions. We did not measure Pel expression and the contribution of Pel to our observations remain to be determined.

The type IV pili are surface appendages that promote biofilm formation, colonization to abiotic and biotic surfaces, and facilitate direct bacterial-host cell attachment [63,64]. PilA, encoded by *pilA,* is the major pilin subunit of the PA type IV pilus and acts as an adhesin [65]. Type IV pili-mediated twitching motility is regulated by multiple different pathways including the alternate sigma factor RpoN, the two-component system PilSR, the secondary global regulator C-di-GMP but also modulated by activity of the ATPase PilB or PilT [44,45,66]. In *Salmonella enterica*, phagocytic function requires the adhesin *fimA*, a major type 1 fimbria that closely resembles type IV pilus found in PA [67]. In PA, it is still not known whether the type IV pilus adhesin (pilus tip) and/or pilus-mediated twitching motility is required to initiate phagocytic uptake. The PilT and PilU retraction ATPases are important for surface attachment to epithelial cells, leading to cytotoxicity [68], and the establishment of corneal infections [69]. It is thus plausible that both adhesion and motility functions contribute to enhancing phagocytic antibacterial function, by increasing the proximity between PA and neutrophils and stabilizing their interactions to initiate phagocytic uptake. Ref to Grana Guttman paper WGS data [30], While non-twitching Persistent isolate 565P lacked a *pilA* gene, 505P encoded for a group I *pilA* allele, described as a distinct allele most commonly found among CF clinical isolates, particularly those recovered from CF children [46,70]. Among the five pilin alleles described in PA, the group I pilin may encode for a shorter pilin associated with a lower hydrophobicity and twitching motility compared to others [46,71]. This genetic variation may contribute to the variability of twitching motility among clinical isolates. Furthermore, Tan et al. showed that group I pilin from PA1244 conferred a fitness survival in a mixed murine pulmonary infection model compared to pilin-deficient PA1244, and that the survival advantage was attributable to its resistance to pulmonary surfactant protein A (SPA)-mediated opsonic phagocytosis [72].

Moreover, results from our *in vivo* mixed infection of 565P and 565P overexpressing PilA further suggest that loss of pilus-mediated twitching motility significantly increases bacterial survival *in vivo* over time, leading to enhanced bacterial colonization and persistence. Similarly, Skurnik et al. reported that transposon mutants in type IV pilus genes enhanced cecal colonization [73]. Findings from our group, as well as others, suggest that loss of pilus is likely an important bacterial mechanism that enhances the establishment of bacterial persistence in the host.

There are several limitations to our study. First, PA phenotypes such as Psl and Pel overproduction and loss of pilus-mediated twitching motility are typical of chronic CF infections. This raises the possibility that the Persistent isolates recovered from our patient cohort, despite being defined as “new-onset” infection, may in fact arise from chronic infections not detected previously. Second, the persistent infection outcome following tobramycin eradication therapy was defined by a PA-positive sputum culture 1 week after the end of treatment. A comparison of PA isolates recovered before and after tobramycin treatment by whole genome sequencing would confirm the persistent nature of these infections. However, Stapleton et al recently reported that 16% of CF children with early infection from this cohort had evidence of mixed strain infections by whole genome sequencing [29]. Third, we limited our studies to four representative clinical PA isolates. Even though they display prototypical phenotypes of our larger cohort collection, a study of a larger number of isolates would enhance the generalizability of our findings. Fourth, our *in vitro* and *in vivo* models do not account for intrinsic neutrophil defects associated with CFTR dysfunction or acquired defects secondary to the inflammatory milieu of the CF airway. Primary murine CF neutrophils or mouse models could be used to examine the contribution of a CF host. Finally, we recognize that our *in vivo* infection model does not capture the dynamic neutrophil-mediated antibacterial functions. Models such as *in vivo* phagocytosis [74], or intravital microscopy would further our understanding of PA-neutrophil interactions [42].

In summary, Persistent isolates display strain-specific and can evade neutrophil-mediated bacterial clearance through several mechanisms, notably resistance to OPK. Loss of twitching motility confers resistance to neutrophil-mediated bacterial clearance and likely contributes to the persistence of PA lung infections. Enhancing PA-neutrophil interactions and neutrophil-mediated bacterial clearance could be a therapeutic strategy to improve the eradication of new-onset PA infections in CF patients.

