## Supplemental figures for "Evasion of neutrophil-mediated bacterial clearance in *Pseudomonas aeruginosa* isolates from new-onset infections in cystic fibrosis children"

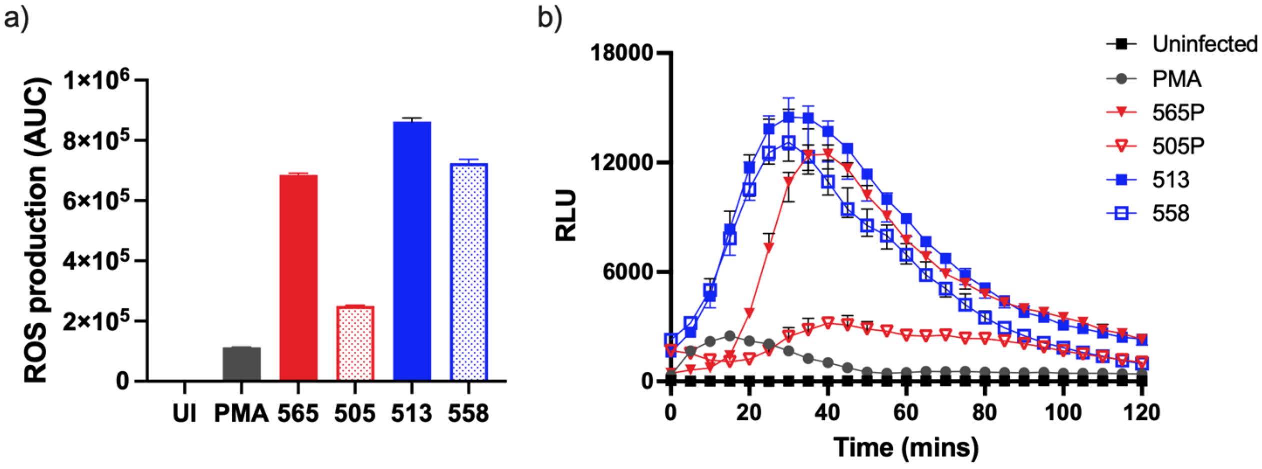


**Figure S1**. **PMA RLU time course.** The area under the curve (AUC) was calculated to determine the relative ROS generated by neutrophils. Results were normalized to ROS induced by phorbol myristate acetate (PMA).

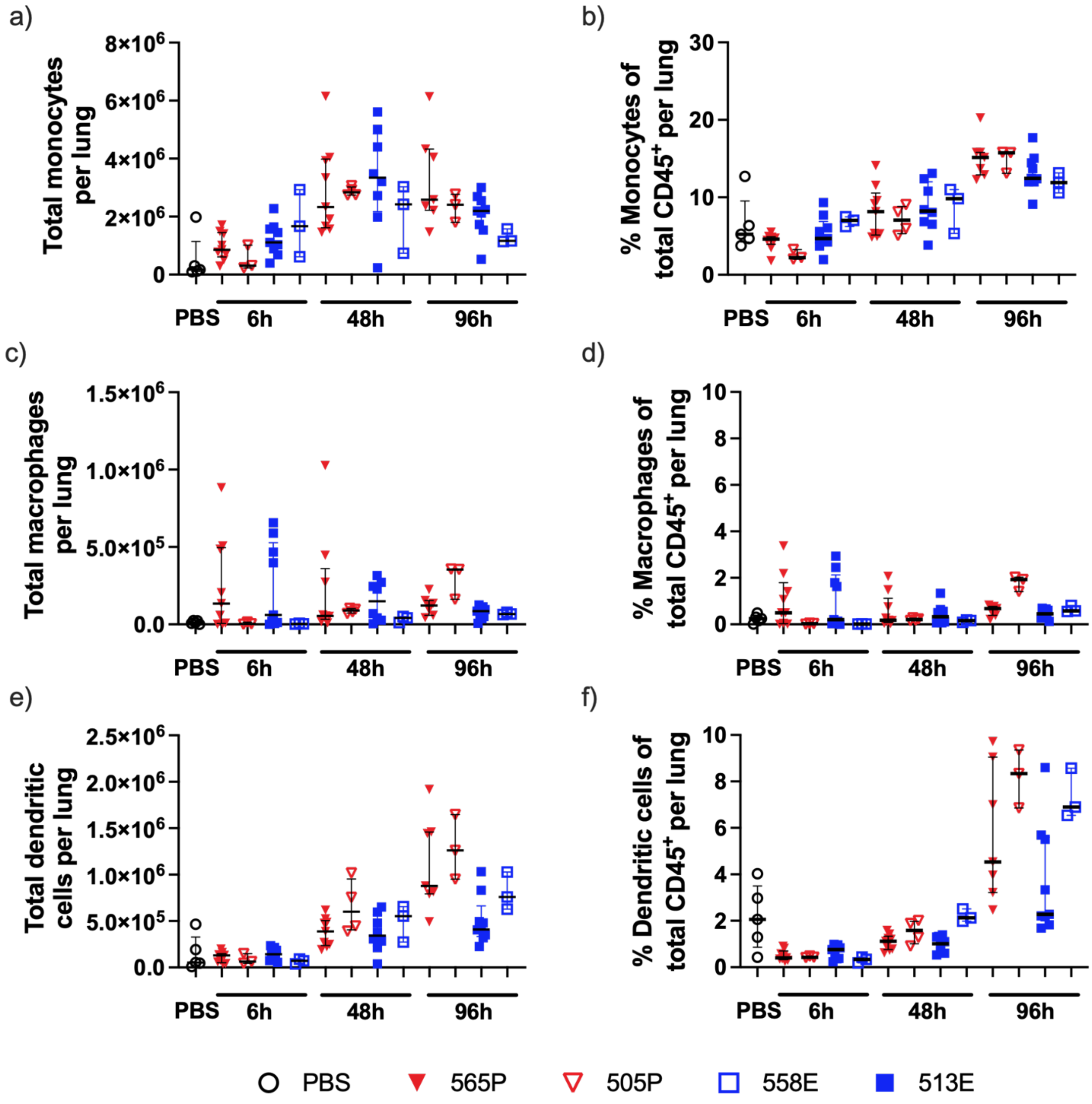


**Figure S2. Immune cell recruitment to the lung in Persistent vs. Eradicated isolate infections.** a) Number of monocytes (CD45-PE, CD11b-APC, Ly6C-BV785), b) Proportion of monocytes, c) Number of macrophages (CD45-PE, CD11b-APC, Ly6G$^{lo}$-BV510, F4/80$^{hi}$-Pacific blue), d) Proportion of macrophages, e) Number of dendritic cells (CD45-PE, CD11b-APC, Ly6G$^{lo}$-BV510, CD11C-BV711), f) Proportion of dendritic cells. Results are shown as median ± IQR. N ≥ 3 mice per strain from ≥ 2 independent experiments. No statistical differences were detected between the groups using ANOVA Dunn's test.

**Figure S3. Comparative genome analysis reveals the absence of *pilA* in 565P contributes to complete loss of twitching.** WGS of PA clinical isolates were used to analyze the sequence of genes linked to pilus-mediated motility (*pilA*, *pilB, fimV, pilT, pilU, pilS, pilR, RpoN, RetS, fliC, algT/U, mucA, mucD, algK, algE, algG, pslA, pslB, pslH, pslI, pslL, pelA, pelB, pelD, pelF, pelG, cdrA, wspA, wspA C, wspA*. Genomic sequence data is available from NCBI BioProject ID: PRJNA893599.

565P (Sample name: Erad565); BioSample: SAMN31427824; SRA: SRS15514966
505P (Sample name: Erad505); BioSample: SAMN31427793; SRA: SRS15514569
513E (Sample name: Erad513); BioSample: SAMN31427802; SRA: SRS15514579
558E (Sample name: Erad558); BioSample: SAMN31427819; SRA: SRS15514599

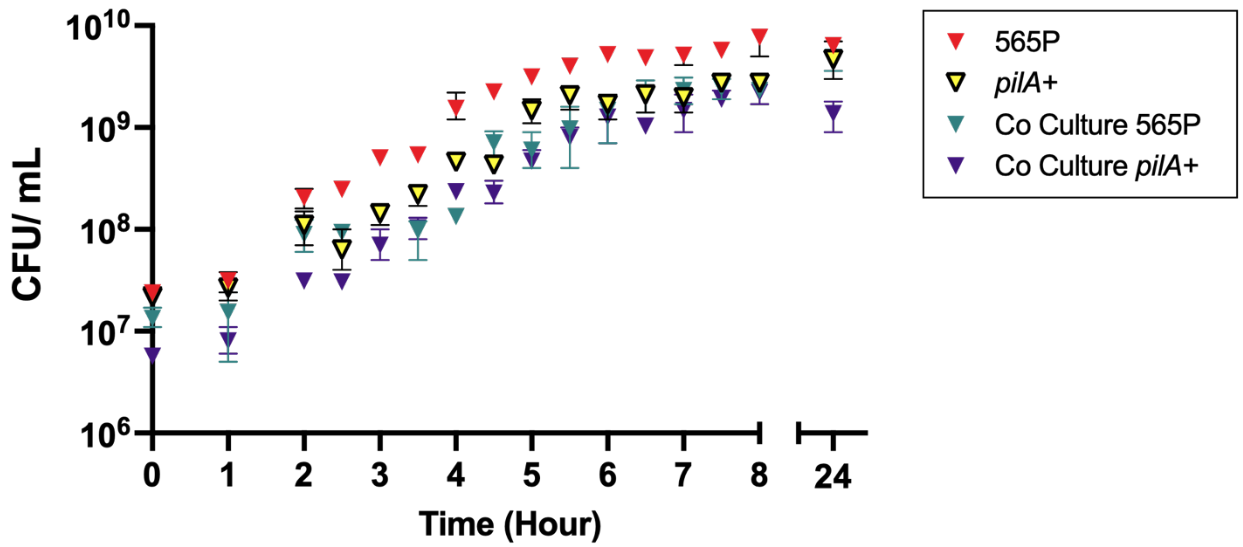


**Figure S4. 565P and *pilA*+ displayed no evidence of *in vitro* inter-strain competition or fitness advantage when grown in planktonic co culture.** 565P and *pilA*+ were grown in co-culture and independently *in vitro,* in nutritive media up to 24h. The constructed *pilA*+ strain did not display any growth defects compared to the 565P parental strain.
